# Messenger RNAs with large numbers of upstream ORFs are translated via leaky scanning and reinitiation in the asexual stages of *Plasmodium falciparum*

**DOI:** 10.1101/823443

**Authors:** Chhaminder Kaur, Mayank Kumar, Swati Patankar

**Author notes:** Amity University of Biotechnology, Amity University, Panvel, Maharashtra 410206, India (Mayank Kumar).

## Abstract

The genome of *Plasmodium falciparum* has one of the most skewed base pair compositions of any eukaryote, with an AT content of 80-90%. As start and stop codons are AT-rich, the probability of finding upstream open reading frames (uORFs) in messenger RNAs (mRNAs) is high and parasite mRNAs have an average of 10 uORFs in their leader sequences. Similar to other eukaryotes, uORFs repress the translation of the downstream gene (dORF) in *P. falciparum*, yet the parasite translation machinery is able to bypass these uORFs and reach the dORF to initiate translation. This can happen by leaky scanning and/or reinitiation.

In this report, we assessed leaky scanning and reinitiation by studying the effect of uORFs on the translation of a dORF, in this case the luciferase reporter gene, and showed that both mechanisms are employed in the asexual blood stages of *P. falciparum*. Furthermore, in addition to codon usage of the uORF, translation of the dORF is governed by the Kozak sequence and length of the uORF, and inter-cistronic distance between the uORF and dORF. Based on these features whole genome data was analyzed to uncover classes of genes that might be regulated by uORFs. This study indicates that leaky scanning and reinitiation appear to be widespread in asexual stages of *P. falciparum*, which may require modifications of existing factors that are involved in translation initiation in addition to novel, parasite-specific proteins.

## Introduction

Malaria affects millions of people in tropical and sub-tropical regions of the world. Over the years, many attempts have been made to control the disease and recently, the World Health Organization renewed the call towards the global eradication of malaria by 2030 (World Malaria Report, 2018). These efforts could be thwarted through resistance to anti-malarial drugs and as a result, new drug targets are still being identified, especially for the most virulent species of the parasite that causes malaria: *P. falciparum*. In this parasite, an essential pathway proposed as a drug target is the protein synthesis machinery (Goodman et al., 2016). *P. falciparum* employs eukaryotic protein synthesis machinery for translation of nuclear-encoded mRNAs, however, an endosymbiotic organelle, the apicoplast, carries out translation using machinery that resembles prokaryotes (Chaubey et al., 2005; Fichera and Roos, 1997; Roy et al., 1999) and is the target of several anti-malarial drugs (Goodman et al., 2016; Pasaje et al., 2016). Interestingly, the cytosolic translation machinery of *P. falciparum* also exhibits differences from that of the human host (Jackson et al., 2011; Wong et al., 2014) and hence, has also been proposed to be a drug target (Sheridan et al., 2018; Wong et al., 2017). Another noteworthy distinction between cytosolic translation of the parasite and its host, is a result of the high AT content (80-90%) of the *P. falciparum* genome. As start and stop codons are AT-rich, this biased sequence leads to the presence of numerous upstream open reading frames (uORFs) in the 5’ leader sequences of messenger RNAs (mRNAs) (Caro et al., 2014; Kumar et al., 2015).

Unlike human and mouse genomes from which ~50% of the transcribed mRNAs would be expected to contain one or more uORFs (Calvo et al., 2009; Ye et al., 2015), ~98% of *P. falciparum* mRNAs are predicted to contain an average of ~10 uORFs per coding sequence (CDS) (Caro et al., 2014; Kumar et al., 2015; Srinivas et al., 2016). The presence of such large numbers of uORFs has serious implications for translation. The scanning model of translation initiation in eukaryotes proposes that the ribosome recognizes the 5’ cap and moves along the transcript until the start codon of the CDS is reached (Aitken and Lorsch, 2012; Hinnebusch et al., 2016; Kozak, 1978). Scanning ribosomes will surely encounter numerous uORFs in the transcripts of *P. falciparum* before reaching downstream ORFs (dORFs), which are the protein coding sequences (CDS) that encode the multitude of proteins required for the parasite’s complex life cycle.

As is the case with other eukaryotes (von Arnim et al., 2014; Child et al., 1999; Hood et al., 2009; Johnstone et al., 2016; McGeachy and Ingolia, 2016), uORFs in *P. falciparum* can down-regulate the expression of the dORF (Amulic et al., 2009; Kumar et al., 2015). This post-transcriptional gene regulation (PTGR) mediated by uORFs takes place by engaging or stalling the ribosome at the uORF, subsequently decreasing the probability of the ribosome initiating translation of the dORF (Morris and Geballe, 2000). It is remarkable that despite the frequent occurrence of uORFs in the majority of transcripts of *P. falciparum*, ribosomes are still able to translate the coding sequences present downstream (Lasonder et al., 2002).

One mechanism through which the ribosome can reach the start codon of the dORF is leaky scanning where the ribosome skips the start codon of the uORF, continues scanning and reaches the AUG of the dORF to start translation (Kozak, 1984; Liu et al., 1984). This strategy is perhaps the most metabolically efficient way of dealing with huge numbers of uORFs. The strength of Kozak sequence flanking the start codon of the uORF has a major role in determining whether that uORF will be skipped (Kozak, 1986). Indeed, it has been reported that the degree of protein expression from the dORF can be regulated by varying the strength of the Kozak sequence of the uORF in mammalian cells (Ferreira et al., 2013). In contrast to other eukaryotic Kozak sequences where the −3 and +4 positions appear to be key, in *P. falciparum* nucleotides preceding the start codon (−5 to −1) do not play a significant role in determining the strength of the Kozak sequences (Kumar et al., 2015). Instead, it seems to be majorly dependent on the nucleotide present at the +4 position.

Another mechanism employed by eukaryotic ribosomes to reach the dORF is reinitiation, where the ribosome translates the uORF and then reinitiates translation at the downstream AUG (Hinnebusch et al., 2016; Hughes et al., 1984; Johnstone et al., 2016; Kozak, 1984). This has been observed for the *var2csa* transcript in *P. falciparum* which harbours a uORF, 360 bases in length, responsible for repression of the *var2csa* coding sequence (Amulic et al., 2009; Bancells and Deitsch, 2013). However, in the presence of a reinitiation factor, *Plasmodium* Translation Enhancing Factor (PTEF), reinitiation is greatly increased at the main ORF leading to synthesis of the VAR2CSA protein (Chan et al., 2017). This phenomenon has clinical significance as expression of VAR2CSA results in adherence of infected red blood cells to ligands in the placenta and manifestation of pregnancy-associated malaria (PAM) (Salanti et al., 2003, 2004). When compared to cultured asexual stage parasites, PTEF (PFB0115W) is highly up-regulated in parasites isolated from the placenta of women suffering from PAM (Francis et al., 2007; Vignali et al., 2011).

Apart from reinitiation factors such as PTEF, translation reinitiation of the dORF in eukaryotes is affected by sequence features of the mRNA leader. Studies in mammalian cells have shown that longer inter-cistronic lengths between the uORF and the dORF increase the probability of reinitiation. It has been proposed that after translating the uORF, the ribosome needs to re-acquire the ternary complex to initiate translation at the next ORF (Kozak, 1987). Therefore, longer inter-cistronic lengths between ORFs would increase the probability of the ribosome acquiring the ternary complex and reinitiating translation (Child et al., 1999; Luukkonen et al., 1995).

Another factor that affects translation of the dORF is the length of the uORF. Translation efficiency of the dORF decreases as the length of the uORF increases (Kozak, 2001; Luukkonen et al., 1995). It is proposed that initiation factors involved in scanning remain bound to the 40S subunit of the ribosome briefly after translation has been initiated. As the ribosome elongates a uORF with a longer length, these factors dissociate from ribosome, thereby decreasing the probability of reinitiation at the dORF. Other structural features of the uORF that affect translation of the dORF are uORF codons and secondary structure of the mRNA in mammalian cells (Hood et al., 2009; Kozak, 2001; Luukkonen et al., 1995).

Considering the large number of uAUGs and uORFs present in *P. falciparum* mRNAs, it is of interest to understand how ribosomes are able to reach the dORF. In this report, we study the stages of the intra-erythrocytic developmental cycle (IDC) to investigate the effect of varying three features viz. Kozak sequence, uORF length and inter-cistronic length on translation of a dORF in *Plasmodium falciparum*. Codon usage is also expected to affect the ability of an uORF to influence translation at the dORF, however, as codon usage tables are available for *P. falciparum* (Nakamura et al., 2000; Saul and Battistutta, 1988) and algorithms for calculating codon adaptability indices (CAI) also well studied (Sharp and Li, 1987), this feature is not assessed in this report. Classes of genes that contain uORFs having each of the features are further studied using gene ontology (GO) enrichment analysis. We also attempt to shed light on the conundrum that the majority of *P. falciparum* transcripts contain multiple uORFs, yet translation of the CDS still takes place. This work brings new insights into the mechanisms of cytoplasmic translation of mRNAs during the asexual life cycle of *P. falciparum*.

## Materials and Methods

### Mutational analysis of the Kozak sequence of the reporter gene

Plasmid Pf86 was treated with BstBI restriction enzyme (Thermo Fisher Scientific) whose recognition sites are present 16 nucleotides upstream and 165 nucleotides downstream of the start codon of luciferase reporter gene. This fragment was replaced by a fragment which had different Kozak sequence. DNA fragments with different Kozak sequences were generated by SDM of plasmid Pf86 using a set of forward primers to introduce desired Kozak sequences and a common reverse primer complementary to ~165 nucleotide region. The primers used for this cloning are included in the Supplementary datasheet 3. The PCR products were then digested with BstBI and ligated to BstBI digested plasmid Pf86. The resulting colonies were screened for insert with desired mutation by restriction digestion as well as colony PCR. Clones were confirmed by sequencing.

### Generating clones for recombinant firefly luciferase, expressed in bacteria

Wild type firefly luciferase had ‘G’ at its +4 position. To generate mutants of the gene with ‘A’/’T’/’C’ at +4 positions, SDM was performed with a forward primer having the desired mutation and a reverse primer complementary to the end of the CDS. Sequence of the primers are included in the Supplementary datasheet 3. This fragment and the vector pET43a were separately digested with XhoI and NdeI restriction enzymes (Thermo Fisher Scientific). The insert fragment was ligated in the digested vector and transformed into *E. coli* DH5α. Colonies were screened by digestion and clones were confirmed by sequencing. The resulting plasmid had a luciferase gene with mutations at +4 position driven by the lac promoter and a C-terminal 6-His tag for purification.

### Induction and purification of variant forms of firefly luciferase

pET43a plasmids containing the firefly luciferase coding sequences with ‘G’, ‘C’ and ‘A’ at +4 positions were transformed into *E. coli* BL21 competent cells. To obtain variants of luciferase protein, the secondary culture of each variant was induced with 0.25 mM Isopropyl β-D-1-thiogalactopyranoside (IPTG). This was followed by 16 hours of incubation at 18°C under shaken condition until the OD_600_ reached 0.6. The harvested cells were re-suspended in 15 ml of ice-cold binding buffer (50 mM NaH_2_PO_4_, 300 mM NaCl and 10 mM Imidazole). The cells were lysed by sonication (Vibrosonics) at 60% amplitude with a pulse of 13 seconds (3 seconds ON and 10 seconds OFF) for 30 minutes under cold condition. Cell lysates were centrifuged at 16000g at 4°C for 15 minutes. The supernatants were loaded onto pre-equilibrated Ni-NTA column (Qiagen Ni-NTA Superflow Cartridge) and the proteins were purified as per the manufacturer’s protocol. The enzymes were eluted in 5 ml of Elution buffer (50 mM NaH_2_PO_4_, 300 mM NaCl and 500 mM Imidazole) and dialysed overnight at 4°C in dialysis tubes (Spectrum Labs Float-A-Lyzer G2 Dialysis Device) overnight in 1.5 litres of dialysis buffer (50 mM NaH_2_PO_4_ and 300 mM NaCl). Purified enzymes before and after dialysis were analysed on SDS-PAGE gel to check the purity of the enzymes.

### Luciferase assay of recombinant luciferase proteins

Protein concentrations of purified luciferase enzyme variants (+4G, +4C, +4A, and +4T) were quantified using the Bicinchoninic Acid (BCA) kit by following manufacturer’s protocol (Sigma-Aldrich). Equal amounts of each luciferase variant was diluted in Passive Lysis Buffer (Promega) and freeze-thawed for three cycles in liquid nitrogen to mimic the conditions of luciferase assay done with parasites. The assay was performed using 90 µL of LAR reagent (Promega) and 10 µL of diluted proteins. Readings were captured for 30 seconds using a luminometer (Berthold Junior LB 9509).

### Removal of the native uORF from the Pf86 plasmid

Plasmid Pf86 was used to test the effect of uORF on the expression of firefly luciferase reporter gene which is flanked by 5’ leader sequence and 3’ UTR of *Pf* heat shock protein 86 (*Pfhsp86*; PF3D7_0708400). In this plasmid, 1837 base pairs (comprising the promoter region and leader sequence of 686 base pairs) from *Pfhsp86* are cloned upstream of the firefly luciferase coding region. The native uORF present 474 bases upstream in the 5’ leader was removed by mutating the start codon (ATG) to TTG by site directed mutagenesis to get Pf86* devoid of any uORF in the 5’ leader. Primers used for this cloning are included in Supplementary datasheet 3.

### Site Directed Mutagenesis (SDM) to introduce uORFs

SDM was used to introduce different uORFs in 5’ leader sequence of *Pfhsp86* cloned in plasmid Pf86*. Non-overlapping forward and reverse primers containing the desired mutation were phosphorylated by polynucleotide kinase (New England Biolabs) as per the manufacturer’s protocol. PCR was carried out by Q5 HiFi DNA polymerase (New England Biolabs). Primers used for SDM are included in Supplementary datasheet 3. Purified PCR product was treated with FastDigest DpnI restriction enzyme (Thermo Fisher Scientific) to eliminate the parental plasmid vector. Linear PCR product was circularised by ligating. The final product was transformed into competent *E. coli* DH5α. The clones were confirmed to contain the desired mutation by sequencing.

### Increasing the length of uORF4 by introducing repeating units

The sequence of uORF4 was designed in way that it contained a recognition site for restriction enzyme AvrII (Thermo Fisher Scientific). The length of this uORF was thus increased by digesting the plasmid uORF4-191 and ligating it to annealed oligonucleotides of increasing length. In this process, the recognition site was lost after ligation. Screening was done by restriction digestion reaction with AvrII and colony PCR to confirm the presence of insert. The sequence of the oligonucleotides used to generate the series of plasmids with increasing uORF4 length is included in the supplementary datasheet 3.

### Parasite culture

*Plasmodium falciparum* 3D7 strain was cultured *in vitro* in media containing RPMI 1640 (Gibco^TM^) supplemented with 0.5% Albumax (Gibco^TM^), 50 mg/L hypoxanthine (Sigma-Aldrich), 2 gm/L D-Glucose (Sigma-Aldrich), 2 gm/L Sodium bicarbonate (Sigma-Aldrich), and 56 mg/L of Gentamycin (Abbott). Culture was maintained at 3% haematocrit using human RBCs obtained from volunteers after approval from the Ethics Committee of IIT Bombay. Whenever required, the parasites were synchronised with 5% D-Sorbitol (Sigma-Aldrich).

### Transfection and luciferase assay

Transient transfection of *P. falciparum* 3D7 was carried out by following the pre-loaded RBCs protocol (Deitsch et al., 2001). Plasmid containing Renilla luciferase gene was used as control to analyse different transfection reactions. 100 µg each of the plasmids—desired plasmid construct with insertion/point mutation and plasmid containing Renilla control—were co-transfected into uninfected RBCs. The transfected RBCs were washed with 10 mL of RPMI media. Late trophozoite stage infected RBCs were added to the preloaded RBCs to give final parasitemia of 0.2 – 0.4%. Media was changed every 24 hours until the luciferase assay was carried out 85 – 90 hours post transfection. The parasite pellet obtained after saponin treatment was re-suspended in 60 µL of 1X Passive Lysis Buffer (Promega). Parasites were lysed by flash freezing in liquid nitrogen and thawing at 37°C for three cycles followed by centrifugation at 10,000 rpm for 2 minutes to remove the cell debris. Luciferase assay was performed using the Dual-Luciferase kit (Promega) as per the manufacturer’s instructions. Relative Light Units (RLUs) were measured over 30 seconds by with a luminometer (Berthold Junior LB 9509).

### Gene Ontology (GO) term enrichment analysis

A Python script was written to extract uORFs from the HMM-defined 5’ leaders of 3137 transcripts sequenced in asexual blood stages. The nucleotide at the +4 position of each uORF, the length of the uORF, and the inter-cistronic length from the CDS was calculated. The list of uORFs with these features is given in the Supplementary datasheet 1. Categorisation of uORFs on the basis of different features (nucleotide at +4 position, length, and inter-cistronic length) was done. GO term enrichment analysis was done by using the GO enrichment tool of PlasmoDB (Aurrecoechea et al., 2009).

### Codon Adaptability Index (CAI) calculations

The CAI of all the uORFs used in this study was calculated by using the algorithm given in Sharp and Li, 1987. The frequency of each codon in *P. falciparum* was obtained from Nakamura et al., 2000.

## Results

### The +4 position of the Kozak sequence plays a major role in translation initiation

Previous work indicated that the +4 position of the Kozak sequence plays a critical role in translation initiation while the nucleotides preceding the start codon (−5 to −1) have no significant contribution towards the strength of the Kozak sequence in *P. falciparum* (Kumar et al., 2015). In this report, the effect of a ‘T’ at the +4 position could not be determined as changing the ‘G’ at the +4 position to ‘T’ in the luciferase expression vector (Pf86), altered the wild type codon ‘GAA’ to a stop codon. To address this lacuna, we decided to change the +5 position to ‘C’ in Pf86 such that now a ‘T at the +4 position would generate a TCA codon.

The plasmid Pf86 was modified to generate seven constructs where the luciferase start codon was surrounded by Kozak sequences all of which had ‘T’ at the +4 position and different sequences from the −5 to −1 positions. These were transiently transfected into *P. falciparum* 3D7 parasites, however, due to low transfection efficiencies, it was not possible to check the mRNA levels of the transcripts. Thus, in a direct comparison to the data reported for ‘A’, ‘C’ and ‘G’ at the +4 position (Kumar et al., 2015), the results shown in this section have been described as the sum of mRNA levels and translation and the word “expression” is used. To determine the effect of different Kozak sequences on expression, reporter gene activity was measured and compared with the wild-type Pf86.

It was observed that despite differences in the −5 to −1 positions, the luciferase activity for each construct with ‘T’ at +4 position was ~60% of the control (Figure 1a). Consistent with the earlier study, the strength of the different Kozak sequences having ‘T’ at the +4 position and varying nucleotides at the −5 to −1 positions, does not correlate with the frequency of each Kozak sequence in the genome of *P. falciparum* (Figure 1a). Additionally, constructs with Kozak sequences having the same nucleotide sequence at the −5 to −1 positions but the four different nucleotides at the +4 position, showed large differences in reporter gene activity (Figure 1b). Here, the luciferase readings for constructs with ‘G’, ‘C’, and ‘A’ at the +4 position in their Kozak sequences have been taken from Kumar et al., 2015 and replotted with construct having ‘T’ at the +4 position in the Kozak sequence. This comparison reinforces the observation that the +4 position is the major determinant of the strength of the Kozak sequence in *P. falciparum*.

**Figure 1:**
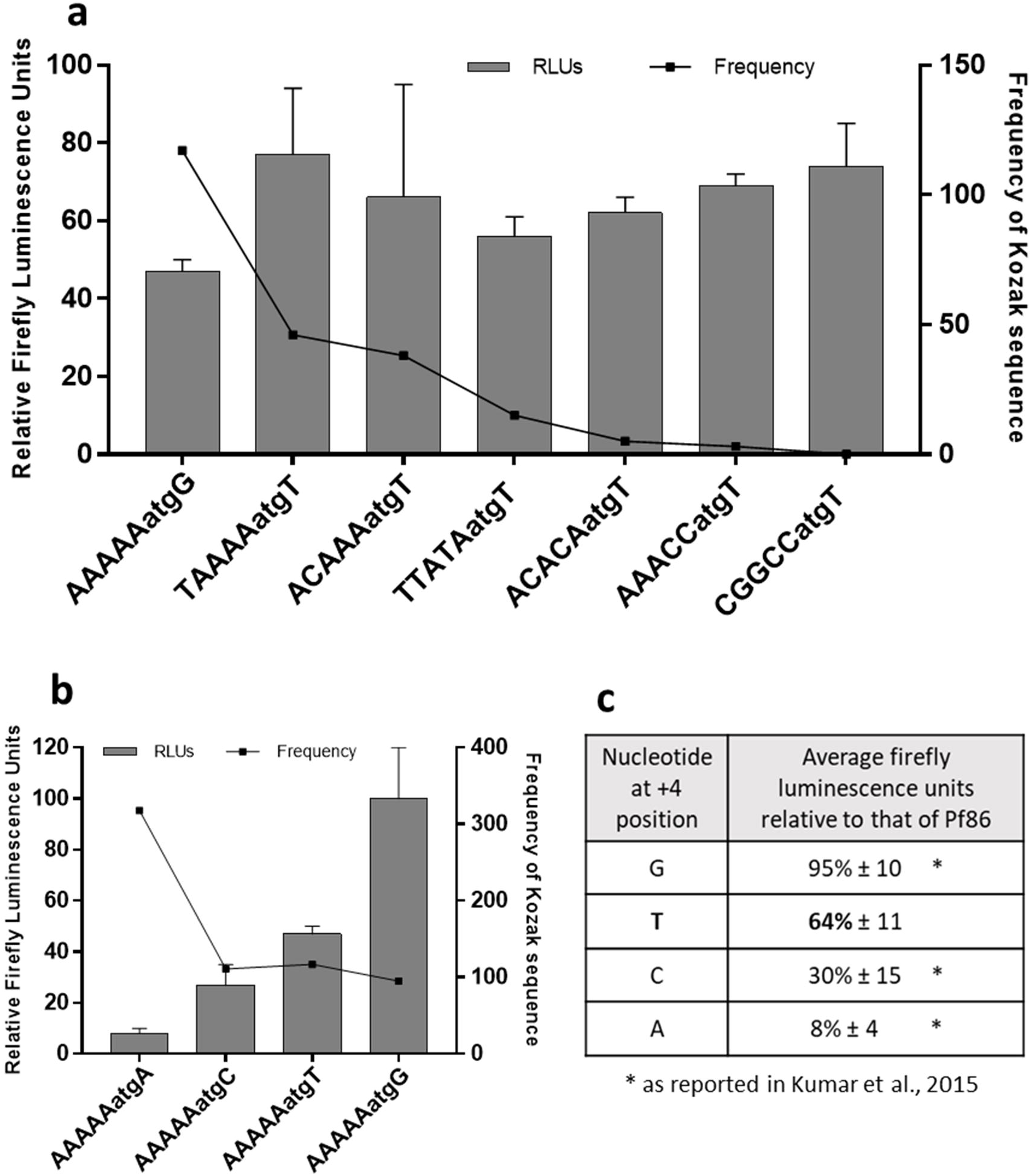
a) Relative firefly luciferase readings of the different constructs of plasmid Pf86 in which the start codon of reporter luciferase is surrounded with Kozak sequences having ‘T’ at +4 position is shown in the bar graph. The frequency of occurrence of the Kozak sequence in the genome of *P. falciparum* is plotted on the secondary Y-axis as a line. Firefly luminescence units were normalized against those of Renilla luciferase for each construct. The firefly luciferase readings of each construct is given with respect to normalized firefly readings of Pf86 with Kozak sequence CGGCCatgG (G at +4 position). Luminescence readings have been measured in Berthold luminometer. Firefly luciferase readings and standard deviations have been calculated from three replicates. b) Firefly luciferase readings (Bar graph) of different constructs of the plasmid Pf86 with Kozak sequence AAAAAatgN, where **N** is A, T, C, or G. The frequency of occurrence of the respective Kozak sequence is shown on the secondary Y-axis as a line. c) Table summarising the average firefly luminescence units obtained from constructs with different nucleotides at +4 position in the Kozak sequence. The reading for G, T, C, and A at +4 position have been calculated by averaging the firefly luminescence units obtained from three, seven, five, and thirteen constructs respectively (Kumar et al., 2015).

It is important to note that changing the +4 position changes the second amino acid of the reporter gene from wild type glutamate (GAA) to glutamine (CAA), lysine (AAA) or serine (TCA). The amino acid differences at the second position might have effects on luciferase reporter activity, as deleting the first seven amino acid residues from firefly luciferase resulted in a significant decrease in the enzyme activity of the protein (Wang et al., 2002). If this was the case for the mutants reported here, the altered luciferase activities would be the outcome of altered luciferase enzyme, rather than alterations in the expression levels of the enzyme due to the Kozak sequence.

In order to eliminate this possibility, the four variants of luciferase gene corresponding to the luciferase proteins generated by making mutations at the +4 were cloned into pET43a expression vector and expressed as recombinant proteins. The activities of these recombinant luciferase enzyme variants were compared to the wild type luciferase enzyme that has glutamate immediately following the start methionine. It was found that changing the second amino acid from glutamate (GAA) to glutamine (CAA), lysine (AAA) or serine (TCA) in the firefly luciferase enzyme did not change the luciferase activity significantly (Single factor ANOVA test; p = 0.1026) (Figure 2). Extrapolating from this data, luciferase enzyme activity in parasites transfected with constructs containing each of the four nucleotides at the +4 position of the Kozak sequence in Pf86, would not result from non-functional enzymes and instead, could be explained by changes in translation initiation driven by altered strengths of the *P. falciparum* Kozak sequence.

**Figure 2:**
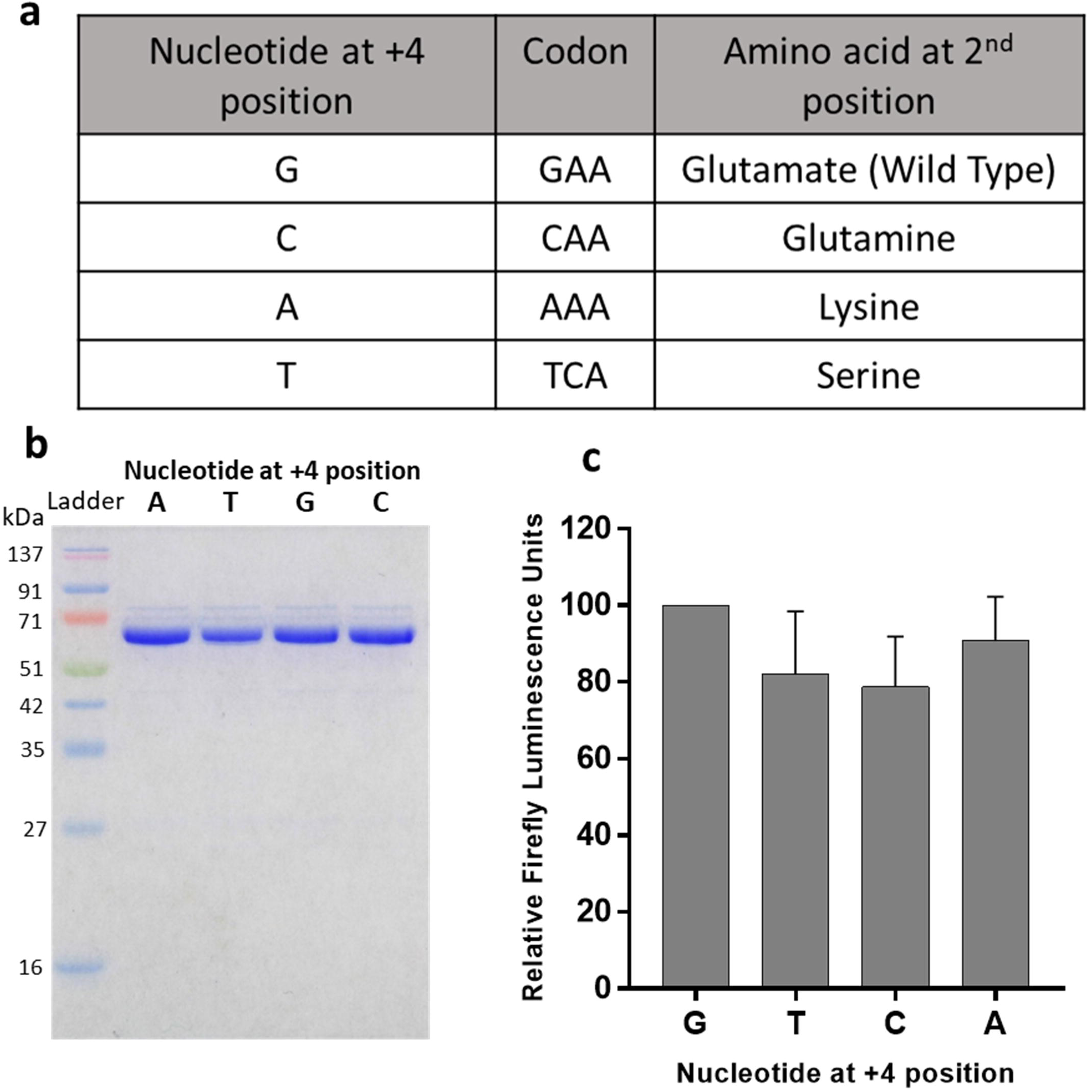
Effect of changing second amino acid of luciferase reporter gene on its activity. a) The table shows the change in second amino acid when wild type ‘G’ at +4 position was changed to ‘C’, ‘T’, or ‘A’. b) Recombinant luciferase enzyme variants with different amino acid at the second position were purified and checked on SOS-PAGE. c) Activity of equal amount of recombinant luciferase variants was tested and relative firefly luminescence units recorded. Single factor ANOVA test; p = 0.1026.

By incorporating the data shown here for ‘T’ at the +4 position with the data published by Kumar and co-workers (Kumar et al., 2015), the average firefly luminescence readings obtained for different Kozak sequences were grouped according to the nucleotide at the +4 position (Figure 1c). Constructs that had Kozak sequences with ‘G’ at the +4 position show the highest reporter gene activity (~100 % as compared with the wild type variant), while constructs containing Kozak sequences with ‘A’ at the +4 position show the lowest activity (~10%) of the reporter gene. The constructs with ‘T’ or ‘C’ at the +4 positions showed intermediate reporter gene activities of ~65% and 30% respectively.

These data led us to assess the +4 positions of coding sequences (CDS) and uORFs in the *P. falciparum* 3D7 genome (PlasmoDB v24), and using bioinformatics, all uORFs were extracted from 5□ leader sequences of 3137 genes expressed in different stages of the IDC (Supplementary datasheet 1). Data for the 5□ leader lengths was exported from a previous study which predicted the length of leader sequences using RNA sequencing of ribosome footprints in RNAs isolated from asexual cultures (Caro et al., 2014).

Greater than 97% of the transcripts analysed contained at least one uORF, as reported earlier (Kumar et al., 2015). A total of 36,769 uORFs were predicted from 3070 transcripts, confirming that there is an average of greater than 10 uORFs per mRNA. The frequency of finding each of the four nucleotides at the +4 position of the Kozak sequences of annotated CDS and uORFs was computed (Supplementary Figure 2a and 2b). This genome-wide analysis revealed that CDS are most likely to have ‘A’ at the +4 position (47%) while ‘G’ follows next, being found in 30% of the CDS. The proportion of CDS with ‘T’ and ‘C’ at their +4 position is 16% and 7% respectively. On the other hand, only 8% of uORFs had ‘G’ at the +4 position, and 9% of uORFs had ‘C’ at the +4 position. As expected, ‘T’ was seen in the +4 position of 48% of the uORFs and ‘A’ was found in 35%, consistent with the AT-bias of intergenic regions (80-90% AT) being higher than that of the CDS (~60-70% AT).

Given that the ‘G’ at the +4 position resulted in a strong Kozak sequence, it is likely that 92% of uORFs have lower probabilities of engaging the ribosome. However, 8% of the total uORFs present in the *P. falciparum* genome have a ‘G’ at the +4 position, indicating that around 2917 uORFs engage the ribosome with high probability and have the potential to repress expression of the downstream CDS. These uORFs may result in lower translation efficiency of the dORFs with which they are associated or may play regulatory roles in translation. Alternatively, although reinitiation is not commonly observed in model eukaryotes, in *P. falciparum*, these data suggest that thousands of uORFs may have properties that allow the ribosome to reinitiate at the dORF.

Reinitiation is suggested by another observation from this analysis. Interestingly, 66% of the 36,769 uORFs predicted in 3137 protein coding transcripts have ‘G’ or ‘T’ at the +4 position, and we show that these types of Kozak sequences engage the ribosome 100% and 60% of the time (Figure 3b). This is consistent with a previous study which reports that approximately half of the total ribosome footprint coverage in 5′ leader sequences of mRNAs overlaps with predicted uORFs (Caro et al., 2014).

**Figure 3.**
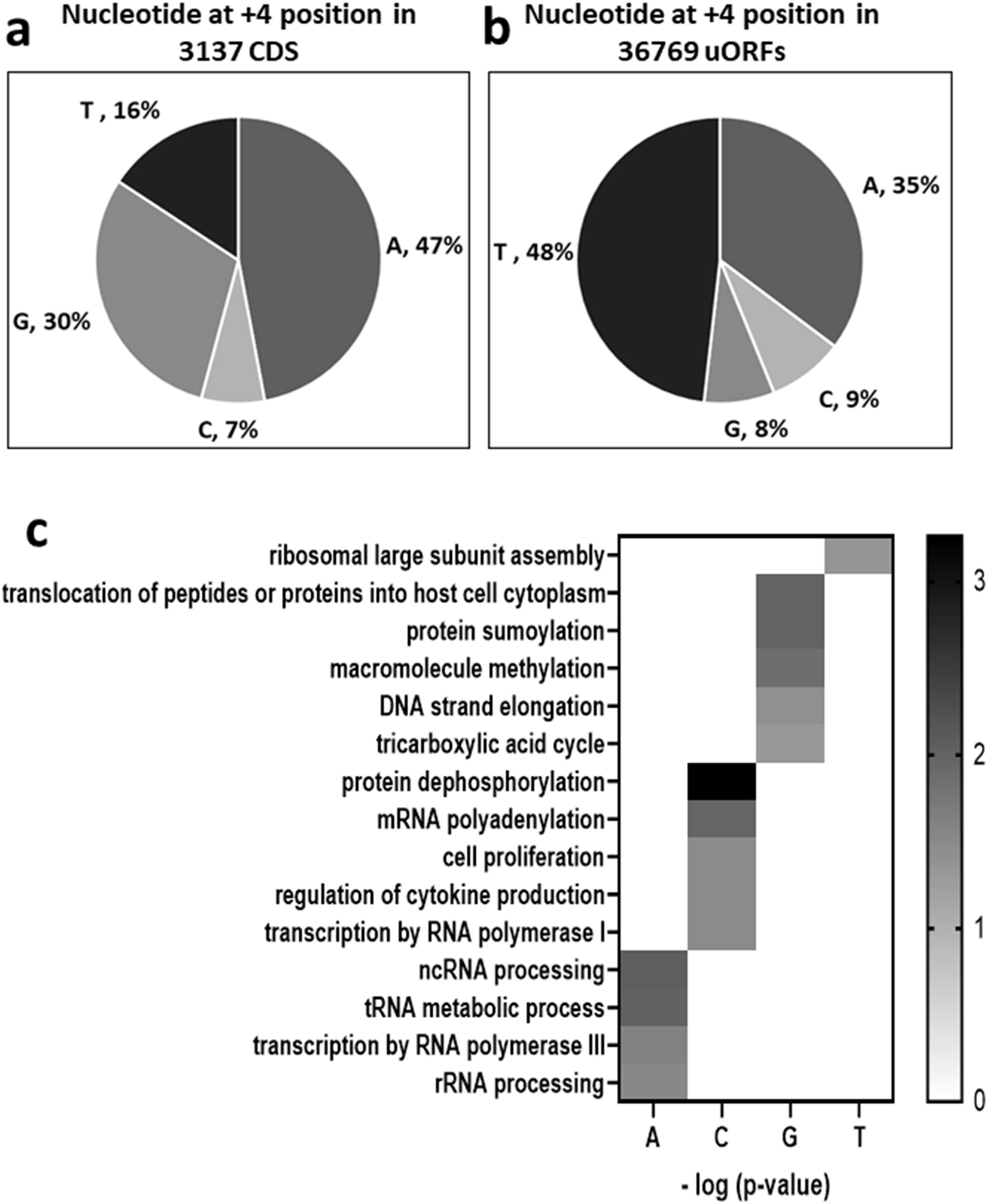
Proportion of four nucleotides (A, T, G, and C) at +4 position in (a) CDS expressed in IDC stages and (b) uORFs predicted in the leader sequence of the CDS. 3137 CDS that are known to be expressed during intra-erythrocytic stage have been taken up for the analysis along with the uORFs that lie in their respective leader sequences (Caro F, 2014) b) Enriched GO terms associated with different nucleotides at +4 position plotted against -log of P-value from Fisher’s exact test (Supplementary datasheet 2). Only the GO terms which are unique for each nucleotide dataset have been plotted in the heat map.

In an attempt to find categories of genes with uORFs having different Kozak sequences, GO term analysis was performed with a p-value cut off of 0.05 using the Gene Ontology Enrichment tool in PlasmoDB (Aurrecoechea et al., 2009). GO terms were enriched for gene sets that are associated with uORFs having Kozak sequences with different nucleotides at their +4 position (Supplementary datasheet 2). The terms that were common between all four sets had high –log (P-value) in the range of 15 and were associated with DNA replication, translation, response to stress, cellular transport, and localization (data not shown). This indicates that uORFs with different nucleotides at +4 position are distributed across different classes of genes.

However, classes of genes having GO terms that were different in each set were seen at lower values of –log (P-value). These classes of genes are shown in the heat map according to their P-value from Fisher’s exact test (Figure 3c). Genes involved in tightly regulated processes such as protein sumoylation and methylation of macromolecules including histones, DNA, tRNA, and proteins, emerged in the set of genes associated with uORFs having ‘G’ at their +4 position. Apart from these broad categories, specific genes such as the translocon component PTEX88/150, involved in translocation of proteins to the host RBC surface, were also seen in this set. Interestingly, an analysis of all CDS in the genome, regardless of whether the gene is transcribed during the IDC, showed a large enrichment of genes involved in host-parasite interactions having ‘G’ at the +4 position of their Kozak sequences. These genes include the *var*, *rifin* and *stevor* genes that play roles in antigenic variation, suggesting that these gene families may be under translational control. Indeed, a member of the *var* gene family, the *var2csa* gene, also has an uORF with a Kozak sequence having ‘G’ at the +4 position and this uORF plays a role translational repression of the dORF (Chan et al., 2017).

Analysis of genes sets associated with uORFs having nucleotide ‘A’ and ‘C’ at their +4 position (weak Kozak context), revealed gene categories that are majorly involved in housekeeping functions. Phosphatases and polyadenylation factors were enriched in the gene set having uORFs with nucleotide ‘C’ at the +4 position. On the other hand, genes that are involved in processing of tRNA, rRNA, and ncRNA were enriched in gene set having uORFs with nucleotide ‘A’ at +4 position. Proteins involved in RNA Polymerase III and RNA Polymerase I transcription were enriched in gene sets with uORFs having ‘A’ and ‘C’ at the +4 position respectively. Since having nucleotides ‘C’ or ‘A’ at the +4 position does not correspond to a strong Kozak sequence, it is likely that the ribosome can scan past these uORFs and reach the dORF by leaky scanning. A list of these uORFs along with their genomic location and dORF has been given in the Supplementary materials.

### Decreasing the inter-cistronic length leads to increased repression of reporter gene expression

The presence of 8% of uORFs having strong Kozak sequences with a ‘G’ at the +4 position suggested that in addition to leaky scanning, the translation machinery of *P. falciparum* might also employ reinitiation to reach the dORF. For reinitiation to occur, after translation termination at the uORF, a fraction of ribosomes would remain associated with the mRNA (Morris and Geballe, 2000). Successful reinitiation at the dORF would depend on the probability of these ribosomes regaining the ternary complex (eIF2, GTP, and initiator Met-tRNA_i_^Met^). In mammalian cells, longer inter-cistronic lengths between the uORF and the dORF lead to efficient reinitiation of the dORF (Kozak, 1987) and similar results have been reported for the *var2csa* gene in *P. falciparum* (Bancells and Deitsch, 2013). It has been proposed that the long inter-cistronic lengths allow the ribosomes to reacquire the ternary complex. It is important to note that inter-cistronic lengths would have no impact on translation of the dORF by leaky scanning, therefore any effect of inter-cistronic length on dORF expression would strongly point towards reinitiation.

Therefore, the effect of changing the inter-cistronic length on expression of the dORF was tested. Here, the 5′ leader sequence of the *hsp86* gene, cloned upstream of a luciferase reporter, was analysed in the plasmid Pf86. The 5′ leader sequence has a native ORF (7 amino acids in length), 474 bases upstream of the luciferase start codon. This uORF was termed native-474 and its translatability score based on the Codon Adaptability Index (CAI) (Sharp and Li, 1987) was computed to be 0.759. This uORF has a ‘T’ at the +4 position of the Kozak sequence, a nucleotide that should give an intermediate Kozak strength (~65% that of ‘G’ at the +4 position).

The start codon of the native ORF was mutated in order to generate a construct which does not have any uAUG/uORF (termed Pf86*). When Pf86* was compared to native-474, it was seen that the native-474 construct gave ~40% of the luciferase activity of Pf86*. Therefore, mutating the start codon of the native ORF led to a 2.5 fold increase in the expression of the reporter gene (Figure 4a). This result is consistent with a previous study which reports that even one uAUG/uORF can repress expression of dORF (Kumar et al., 2015). To test the effect of changing inter-cistronic distance, the native uORF was moved to a position corresponding to 29 bases upstream of the luciferase start codon (native-29). Decreasing the inter-cistronic length to 29 nucleotides gave ~20% luciferase expression (Figure 4a) compared to Pf86*.

**Figure 4.**
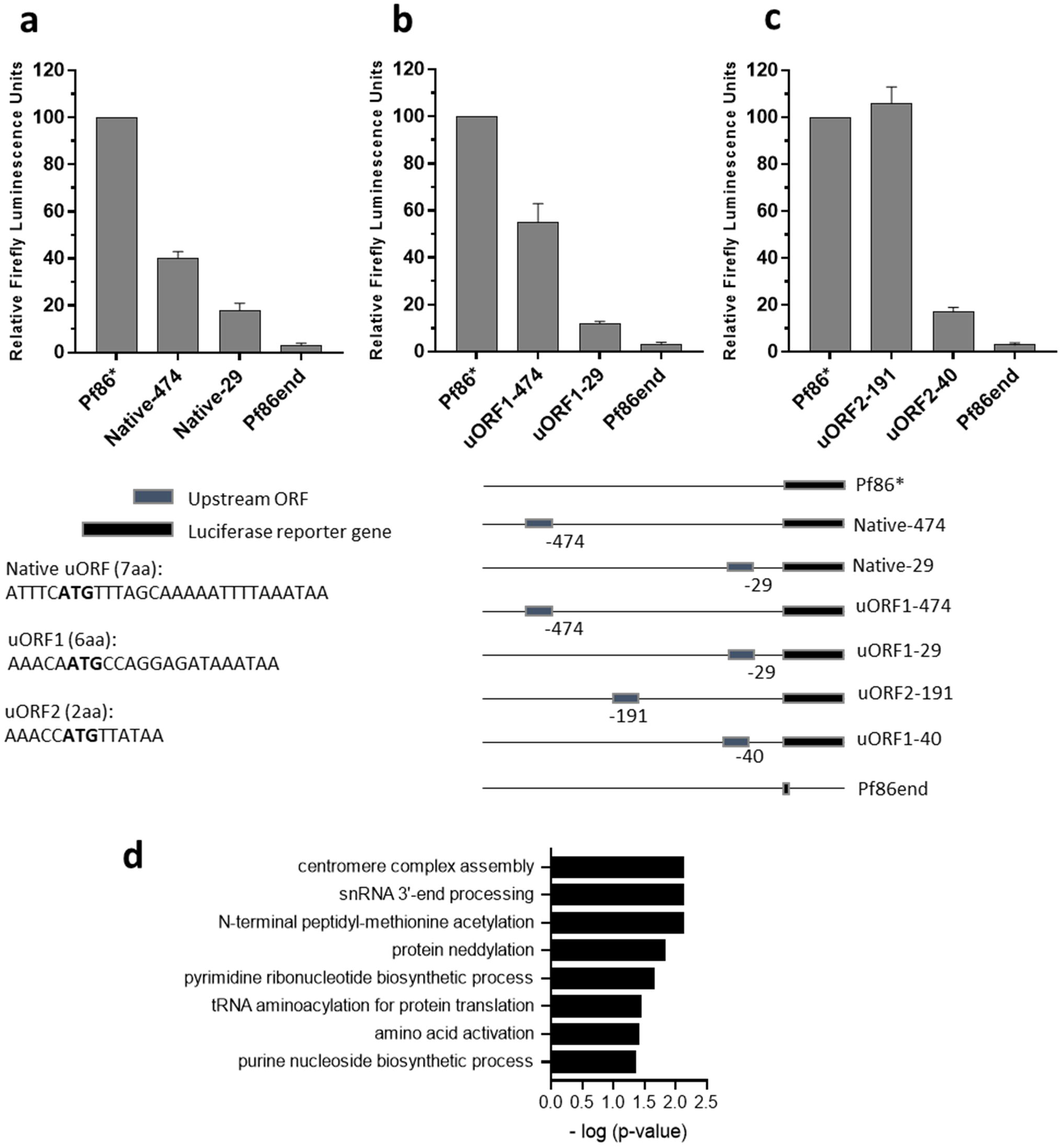
Effect of inter-cistronic distance on the repression of reporter gene expression. Pf86*, a construct which do not have any upstream ORF has been used as positive control while Pf86end, a construct in which the second codon of luciferase reporter gene is mutated to stop codon has been used as negative control. a) Firefly luciferase units obtained when an ORF present natively (Native-474) in the plasmid Pf86 at 474 nucleotides upstream of the start codon was cloned 29 nucleotides upstream of the luciferase reporter gene(Native-29) b) Firefly luminescence units obtained when uORF1 was cloned at 474 and 29 nucleotides upstream of the start codon (uORF1-474 and uORF1-29 respectively) c) Firefly luminescence units obtained when uORF2 was cloned at 191 and 40 nucleotides upstream of the start codon (uORF2-191 and uORF2-40 respectively) d) GO terms enriched in the gene set associated with uORFs with inter-cistronic length less than or equal to 50 nucleotides. The enriched GO terms are plotted against their -log of P-value from Fisher’s exact test (Supplementary datasheet 2)

Another uORF, uORF1 (CAI: 1), was introduced at either 474 or 29 nucleotides upstream of the start codon of luciferase. For these constructs, uORF1-474 and uORF1-29 respectively, luciferase expression reduced to 55% and 12% of the Pf86* control (Figure 4b). The +4 position for the Kozak sequence of uORF1 was ‘C’, a nucleotide that shows weaker ability to engage the ribosome (~30% that of ‘G’ at the +4 position). Both the uORFs tested, native uORF and uORF1, showed approximately equal levels of repression suggesting that repression depends on a combination of Kozak sequences and CAI scores.

Finally, a short upstream ORF, uORF2 (CAI: 1), coding for two amino acids and having a ‘T’ at the +4 position of the Kozak sequence, was introduced in the 5’ leader. Despite having a Kozak sequence of a strength corresponding to 65% of that of the strongest Kozak tested, introduction of uORF2 191 nucleotides upstream of the start codon (uORF2-191) did not reduce luciferase expression. The combination of a strong Kozak sequence with a short coding sequence of merely two amino acids appears to make this uORF one that allows the ribosome to reinitiate at the luciferase gene. This hypothesis will be addressed in a subsequent section of this report. Interestingly, when uORF2 was introduced 40 nucleotides upstream of the luciferase start codon (uORF2-40), luciferase expression reduced to 16% as compared to Pf86* (Figure 4c). In sum, data from all three uORFs tested so far indicates that inter-cistronic length plays an important role in expression of downstream ORF.

From our data, it can be concluded that introducing a uORF close to the start codon of dORF leads to a significant decrease in the expression of the dORF. In all the cases, the maximal reduction in reporter activity was seen when the uORF was introduced within 50 nucleotides upstream of the start codon of luciferase, irrespective of differences in CAI, Kozak sequence and length of the different uORFs. This is similar to mammalian cells, where decrease in the inter-cistronic length leads to inefficient translation of the dORF (Kozak, 1987; Luukkonen et al., 1995). Importantly, as inter-cistronic length has no effect on leaky scanning, the data are suggestive of reinitiation being the mechanism for translation of dORFs in *P. falciparum* when uORFs can engage the scanning ribosomes.

Driven by the experimental results of inter-cistronic lengths, a bioinformatics analysis of the genome of *P. falciparum* was undertaken to extract uORFs that are present at different inter-cistronic lengths from the annotated CDS. Particularly, those uORFs that lie within 50 bases of the CDS were identified as they might cause repression. Using the 5’ leader sequence, we mapped all the uORFs found in the leader sequence all the transcribed CDS in asexual blood stages of *P. falciparum* (Supplementary datasheet 1). GO term analysis of gene sets having uORFs within 50 nucleotides upstream of the start codon showed enrichment of genes associated with centromere and kinetochore assembly, snRNA processing, protein neddylation and ubiquitination, tRNA charging and amino acid activation, and biosynthetic pathways for purine and pyrimidine biosynthesis (Figure 4d)(Supplementary datasheet 2).

Analysis of the uORFs associated with all CDS (regardless of whether they are transcribed in the IDC stages), revealed enrichment of GO terms associated with *var* and *rifin* genes (data not shown). It is well known that *var* genes are subjected to transcriptional control such that out of the repertoire of *var* genes, only one is translated in the late asexual stages (Dzikowski et al., 2007; Kyes et al., 2003; Scherf et al., 1998). Our data on Kozak sequences and inter-cistronic lengths of uORFs associated with *var* genes suggest that for the transcribed *var* genes, there appears to be a high likelihood of translation repression by uORFs which would be relieved by mechanisms such as reinitiation.

### Expression of the reporter gene depends on the length of uORF

So far, results shown in this report have indicated that in asexual stages of *P. falciparum*, reinitiation takes place to allow the ribosome to handle the large number of uORFs found in mRNAs. Reinitiation efficiency depends on the length of the uORF, with longer uORFs presumably resulting in the loss of initiation factors from the ribosome, in turn, decreasing reinitiation efficiency in mammalian cells (Hood et al., 2009; Kozak, 2001; Luukkonen et al., 1995).

Possible effects of the length of the uORF were investigated in a bi-cistronic transcript by introducing ORFs of varying lengths upstream of the luciferase reporter gene. A 12 nucleotide long upstream ORF, uORF4 (CAI: 0.484) was introduced 191 nucleotides upstream of the start codon of the reporter gene. Keeping the Kozak sequence and inter-cistronic distance constant, the length of this uORF was increased by inserting a repeating unit 24 nucleotides in length in a restriction site present in the uORF to give uORFs of different lengths. This cloning strategy generated constructs termed uORF4-1X to uORF4-5X that were 36, 60, 84, 108, and 132 nucleotides long and encoded peptides consisting of 11, 19, 27, 35, and 43 amino acids with CAI values of 0.837, 0.890, 0.911, 0.922, and 0.929 respectively. Plasmid Pf86* with no uORF was used as a control for measuring luciferase reporter activity in transient transfection assays (Figure 5).

**Figure 5:**
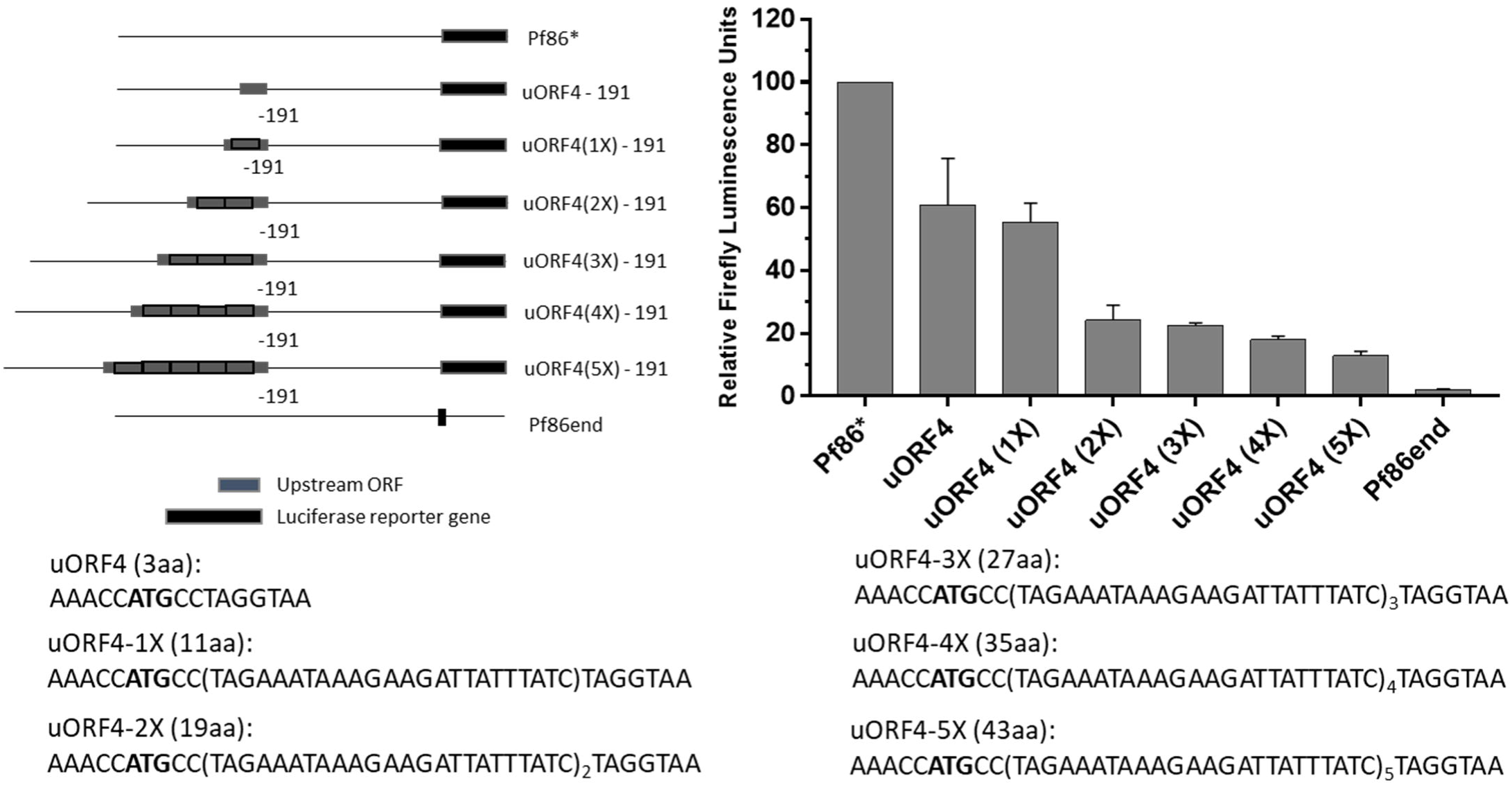
Length of upstream ORF affects the expression of the reporter gene. Pf86* and Pf86end have been taken as the positive and the negative control respectively. Firefly luciferase units obtained when upstream ORFs of different length and same Kozak sequence are cloned at the same distance upstream of the start codon of the reporter gene. Firefly luminescence units obtained in presence of uORF4 and its derivatives with increasing length, uORF4-1X, u uORF4-2X, uORF4-3X, uORF4-4X, and uORF4-5X, upstream of luciferase reporter

Increasing the length of upstream ORF, while keeping the Kozak sequence and inter-cistronic length constant, led to a gradual decrease in reporter gene expression. Although the CAI value of uORF4 was approximately half that of the other uORFs, the CAI values of uORF4-1X through uORF4-5X increased incrementally from 0.837 to 0.929. It is expected that increased CAI values would reflect uORFs that are easier to translate, and should result in a higher probability of the ribosome reaching the dORF and therefore, an increase in luciferase activity. Instead, luciferase activity showed a decrease in the presence of longer uORFs.

This suggests that as in other eukaryotes, in *P. falciparum*, apart from the Kozak sequence and inter-cistronic length, the length of the uORF also has an important role to play in expression of dORF. In the presence of longer uORFs, the ribosomes have a lower probability of reaching the reporter gene and hence, expression is repressed. In accordance with data presented in this report, these results also suggest that reinitiation occurs in *P. falciparum*.

Here we show that uORFs coding for peptides longer than 19 amino acids (60 nucleotides) show repression greater than 50% as compared to the control plasmid having no uORF. As the *P. falciparum* genome has a large number of uORFs, for the dORF to be translated, one would expect that the majority of uORFs should be less than 60 nucleotides long. As expected the average length of predicted uORFs in the transcripts is 45 nucleotides (14 amino acids). The frequency distribution of the lengths of uORFs shows that 67% of ORFs are encoded by sequences less than 45 nucleotides (Figure 6a). The majority of these uORFs would allow the ribosome to reinitiate successfully at the start codon of the CDS. However, longer uORFs (length > 45 nucleotides) may engage the ribosomes for a longer time and hence, reduce efficient reinitiation at the start codon of the dORF. A list of these uORFs and their downstream CDS has been given in the Supplementary datasheet 1.

**Figure 6:**
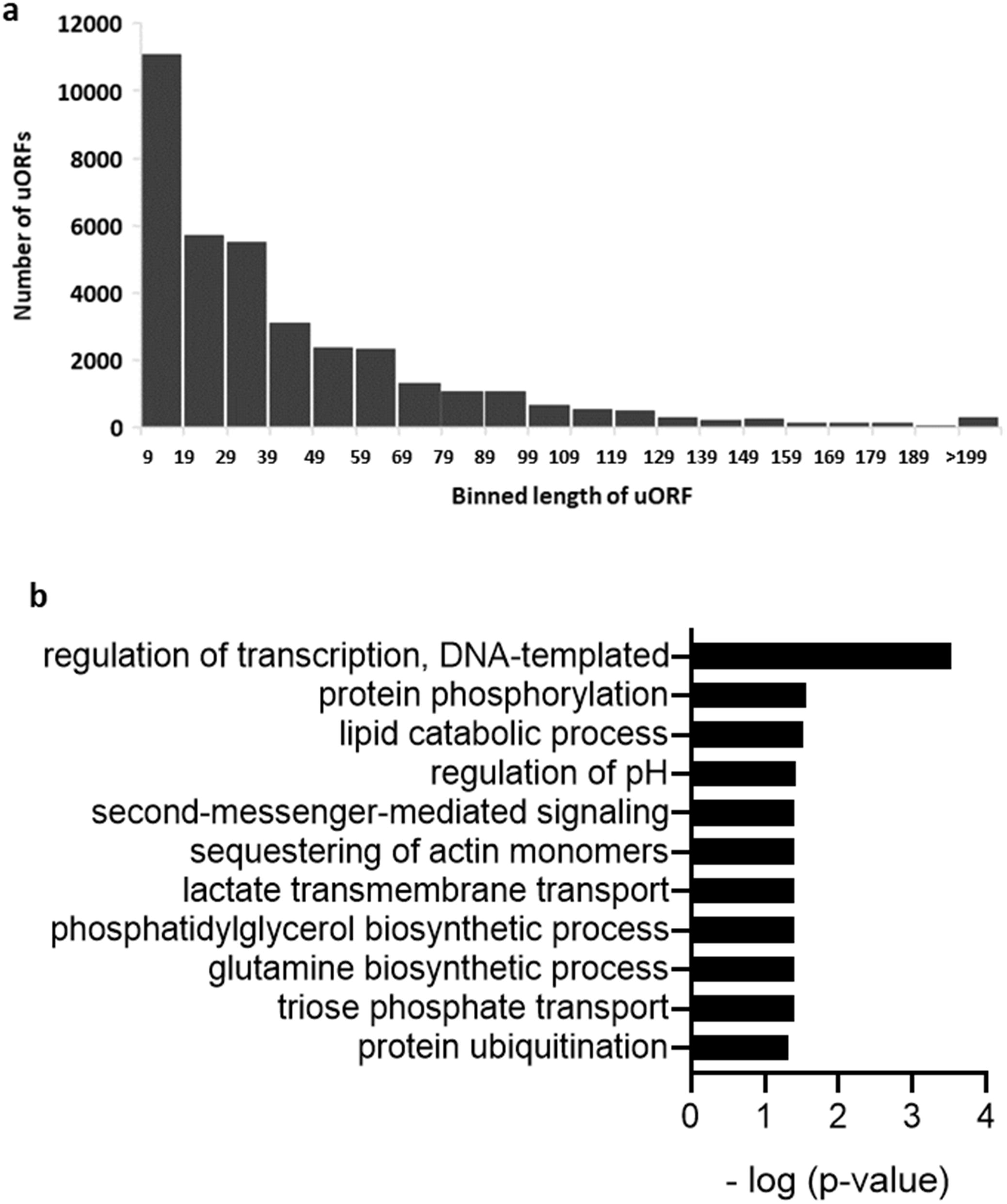
a) Frequency distribution of length of uORFs predicted from 3137 expressed CDS b) GO terms enriched in the gene set associated with uORFs whose length is greater than 199 nucleotides. GO terms are plotted against - log (p­ value) from Fisher’s exact test.

Differential GO term analysis between the sets of genes associated with uORFs whose length is less than and those greater than 45 does not reveal enrichment of any category (data not shown). However, a set of genes that are associated with uORFs with length greater than 200 nucleotides was enriched in GO categories involved in transcription regulation, housekeeping processes (lipid catabolism and pH regulation) and protein ubiquitination and phosphorylation (Figure 6b) (Supplementary datasheet 2). Interestingly, the AP2 domain transcription factor family which is involved in transcriptional regulation during development of *P. falciparum* (Painter et al., 2011) has numerous uORFs that are longer than 200 nucleotides. The uORFs present in the 5□ leaders of these genes might be involved in regulating the expression of this transcription factor and hence, controlling development in the parasite.

### Assessing the contribution of leaky scanning and re-initiation in the expression of a single dORF

Results shown so far have indicated that *P. falciparum* parasites, despite having multiple uAUGs/uORFs in the 5□ leaders of their mRNAs, are able to translate their CDS by leaky scanning to bypass weak Kozak sequences and by reinitiation at the dORF. Reinitiation has been described previously in *P. falciparum* for translation of the *var2csa* gene in the presence of a 360 nucleotide long uORF (Chan et al., 2017), with a parasite translation factor, PTEF, being induced in parasites that are found in the placenta of pregnant women having malaria. Our data show that reinitiation may occur frequently in asexual parasites as well. Next, we analyzed whether leaky scanning and reinitiation occur simultaneously in the asexual stages, in the presence of a single repressive uORF.

The strategy used was to clone a single uORF, out of frame with the dORF (luciferase reporter gene). The next step was to eliminate all the in-frame stop codons downstream of the uORF by mutation. To test the extent of leaky scanning and reinitiation, the stop codon of the uORF is mutated. In case ribosomes rely on reinitiation alone as the mode of translation of the dORF, the expression of the dORF should be eliminated after mutating the uORF stop codon. This is because the uORF is out of frame with the dORF and if initiation takes place, with no in-frame stop codons, the resulting protein will not encode luciferase. In case the ribosomes rely on leaky scanning alone, the expression of the dORF should remain the same as it was before mutating the stop codon. However, if a mix of leaky-scanning and reinitiation occurs, the expression should be less than the expression seen before mutation of the stop codon of the uORF (Figure 7a).

**Figure 7:**
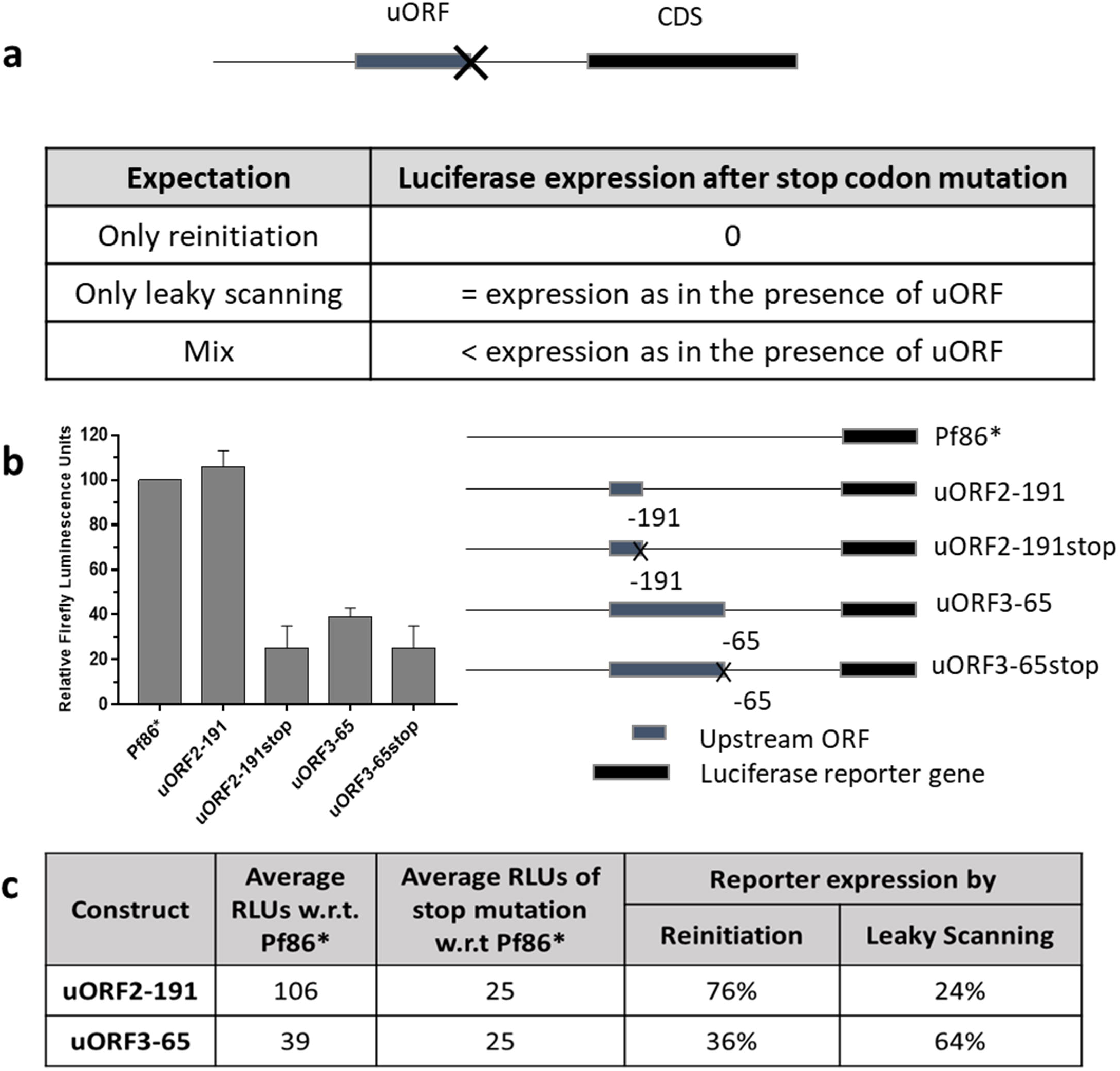
Determining the contribution of reinitiation and leaky scanning in the expression of the reporter gene by mutating the stop codon of a uORF, thereby eliminating the share of reinitiation. a) Schematic of expectations if the expression of the luciferase reporter gene was through only reinitiation, or only leaky scanning, or mix of both reinitiation and leaky scanning. b) Firefly luciferase readings obtained for constructs containing uORFs, uORF2-191 and uORF3-65, present 191 and 65 nucleotides upstream of the luciferase start codon respectively. Mutating all the in-frame stop codons eliminates the chance of reinitiation, leading to the expression of luciferase only via leaky scanning, assuming only these two phenomenon contribute towards expression of the reporter gene. c) Contribution of reinitiation and leaky scanning for two constructs containing uORFs of different lengths and at different inter-cistronic distances. The percentage share for reinitiation and leaky scanning have been calculated by normalising the RLU values.

To assess the contribution of reinitiation in translation of the dORF the construct containing uORF2-191, which does not repress the dORF (Figure 4c) was selected; this uORF allows the ribosome to reach the dORF 100% of the time, providing an excellent scenario to test the contribution of leaky scanning and reinitiation in translation of the dORF. The stop codon of the uORF and all the in-frame stop codons were mutated to create a construct termed uORF2-191stop. It was observed that the expression of luciferase decreased to 25% as compared to Pf86* (Figure 7b). Therefore, 75% of reporter gene expression appears to be dependent on reinitiation and 25% on leaky scanning or other mechanisms. (Figure 7c). In case of uORF2-191, the chances of reinitiating the dORF could be high due to 191 nucleotides in the inter-cistronic length and a small uORF length, paving the way for the ribosome to reacquire initiation factors.

Another uORF, uORF3-65 which is 135 nucleotides in length was introduced 65 nucleotides upstream of the luciferase start codon. The Kozak sequence of uORF2-191 and uORF3-65 are the same. As discussed in the previous sections, the length of this uORF and the short inter-cistronic distance would predict that the ribosomes have a lower probability of reacquiring initiation factors. Therefore, the contribution of reinitiation to the translation of the dORF should be less than that seen for uORF2-191. Consistent with this prediction, the luciferase expression obtained in presence of this uORF was 39% that of Pf86* (Figure 7b). Mutation of the stop codon of uORF3-65 led to ~1.5 times decrease in expression as compared to the case when the stop codon was present. The proportion of leaky scanning and reinitiation were calculated using the same strategy as used for uORF2-191 and seen to be 64% and 36% respectively.

As in the previous sections, the start codon of the uORF was mutated to remove the uORF. The luciferase expression from both constructs uORF2-191ATGmut and uORF3-65ATGmut is restored to the control levels (Supplementary Figure 1), indicating that the experiment reflects translation effects rather than effects on mRNA stability or secondary structures. Therefore, for two uORFs, a mix of leaky scanning and reinitiation is seen for translation of the dORF. These results reinforce the observations that reinitiation may be widespread in asexual stages of *P. falciparum*, and not restricted to parasites isolated from pregnancy associated malaria (PAM) samples.

## Discussion

In this report, a systematic study of the features that contribute to repression by uORFs resulted in the observation that *P. falciparum* asexual stage parasites employ widespread leaky scanning and reinitiation to allow the scanning ribosome to reach the dORF and express a multitude of proteins. Some classes of proteins may undergo translational regulation as uORFs associated with their CDS have features associated with repression: strong Kozak sequences, inter-cistronic lengths less than 50 nucleotides and/or lengths greater than 200 nucleotides. Specifically, these classes of proteins include antigenic variation gene families, including *var*, *rifin* and *stevor*, proteins involved in ubiquitination and members of the AP2 transcription factor family. Potential mechanisms by which translational regulation of these classes of proteins by uORFs might occur are discussed.

### Reinitiation of the downstream ORF is a strategy used by *P. falciparum* to handle large numbers of repressive uORFs

We show that uORF length and inter-cistronic distance affect expression of the downstream luciferase reporter gene, both suggesting that reinitiation takes place in asexual stages of *P. falciparum*. Data suggest that the ribosome is able to reinitiate translation of CDS when uORFs are less than 60 nucleotides long. This is comparable to other eukaryotes where uORFs that allow reinitiation are usually less than 30, 48, and 90 nucleotides in yeast, plants, and mammals respectively (von Arnim et al., 2014; Calvo et al., 2009; Kozak, 2001). In case of inter-cistronic length, we show that uORFs within 50 nucleotides of the start codon repress the expression of dORF drastically in *P. falciparum*. A similar observation has been made for viruses, where expression of the dORF decreases when the inter-cistronic length is reduced from 64 to 16 nucleotides (Luukkonen et al., 1995). Our analysis shows that 40% transcripts in the IDC stages of *P. falciparum* have at least one uORF within 50 bases upstream of start codon of CDS. We propose that to translate the CDS, *P. falciparum* would utilize molecular factors to enhance the probability of reinitiation.

One such player is PTEF (*Plasmodium* translation enhancing factor) which is required for reinitiation of translation of the *var2csa* CDS in the presence of a 360 nucleotide long uORF (Chan et al., 2017). PTEF is expressed in intra-erythrocytic stages with expression levels increasing from ring to schizont stage (Pease et al., 2013). In case of pregnancy-associated malaria, 7-15 fold higher expression of PTEF appears to be needed for translation of *var2csa* due to the presence of a uORF which is significantly longer than the average length of predicted uORFs (Chan et al., 2017). One might speculate that in asexual stage parasites, low levels of PTEF are sufficient to enable reinitiation in the presence of numerous smaller uORFs which are abundant in transcripts. The presence of wide-spread reinitiation, possibly driven by PTEF, leads to the speculation that as this protein shows low sequence identity with human proteins, it might be a potential drug target for the blood stages of *P. falciparum*.

Another molecular player that is involved in reinitiation is the translation initiation factor, eIF2α. Evidence of reinitiation mediated by phosphorylation of eIF2α during nutritional stress responses is well documented for the *S. cerevisiae* GCN4 transcript (Hinnebusch, 1993, 2005) and for the integrated stress response in mammalian cells (Young et al., 2015). In case of *P. falciparum*, one would expect that if there is a significant amount of reinitiation, PfeIF2α should be phosphorylated. Interestingly, phosphorylation of PfeIF2α is not seen in ring and trophozoite stages, however, a sudden increase is observed in schizonts (Zhang et al., 2017). Although suggestive of reinitiation occurring predominantly in schizonts, due to the lag is observed between the peak of mRNA synthesis and corresponding protein abundance in *P. falciparum* (Foth et al., 2011; Le Roch et al., 2004), it is also possible that the phosphorylated PfeIF2α might promote reinitiation from mRNAs expressed at earlier IDC stages.

Another molecular factor that plays a role in reinitiation is eIF3, warranting further work on this protein in *P. falciparum*. Eukaryotic IF3 (eIF3) remains transiently attached to the elongating ribosome and stabilizes the post-termination complex to stimulate reinitiation of the dORF in *S. cerevisiae* (Cuchalová et al., 2010; Hronová et al., 2017; Mohammad et al., 2017). Similarly, the H subunit of eIF3 helps in efficient resumption of scanning after translation of the uORF in *Arabidopsis* (Roy et al., 2010).

### The Kozak sequence determines the extent of leaky scanning

Alleviation of translational repression due to numerous uORFs can also be achieved through leaky scanning, possibly the most metabolically efficient way to handle multiple uORFs. A known contributor to leaky scanning is the strength of the Kozak sequence (Ferreira et al., 2013, 2014; Kozak, 1999). This report confirms published observations that the nucleotide following the start codon (+4 position) plays a significant role in determining the strength of the Kozak sequence in *P. falciparum* (Kumar et al., 2015), unlike other eukaryotes where nucleotides at −3 and +4 positions are both important (Pisarev et al., 2006).

In addition to Kozak sequences, eIF1, a translation initiation factor in the preinitiation complex can facilitate selection of the start codon (Cheung et al., 2007) by release of eIF1 from the preinitiation complex leading to translation initiation (Nanda et al., 2009; Passmore et al., 2007; Pestova et al., 1998). High concentrations of eIF1 lead to stringent selection of AUGs having strong Kozak sequences (Andreev et al., 2015; Fijalkowska et al., 2017; Loughran et al., 2012) and the phosphorylation state of eIF1 under stress conditions also determines the selection of the start codon, helping to bypass start codons with weak Kozak sequences (Zach et al., 2014). Another factor, the m7G-cap binding factor eIF4G1, also enhances leaky scanning of uORFs near the cap when bound to eIF1 in mammalian cells (Haimov et al., 2018). Another factor that affects leaky scanning in eukaryotes is eIF2α. When eIF2α is phosphorylated under stress conditions, uORFs with weak Kozak sequences are more likely to be bypassed by leaky scanning (Palam et al., 2011). Interestingly, eIF2α phosphorylation also facilitates reinitiation (Hinnebusch, 2005), suggesting that this protein could be a key player in handling repression by uORFs.

### Implications of uORFs in gene regulation

Like other eukaryotes*, P. falciparum* utilises a spectrum of mechanisms of gene regulation ranging from epigenetic (Saraf et al., 2016) to post-transcriptional gene regulation (Bunnik et al., 2013; Caro et al., 2014). However, unlike other eukaryotes, *P. falciparum* seems to lack the variety of canonical eukaryotic transcription factors or stage-specific transcription factors (Coulson et al., 2004; Gardner et al., 2002). Stage-specific regulation at the transcriptional level is achieved via the AP2 family of transcription factors (Balaji et al., 2005). This family of transcription factors is known to control development of the parasite through various stages (Painter et al., 2011). Interestingly, transcript and protein levels during each stage of the parasite’s lifecycle are poorly correlated which points towards translationally controlled expression (Le Roch et al., 2004) and a translational regulator, PfALBA1, represses translation of transcripts involved in invasion until the parasite is mature enough to invade (Vembar et al., 2015, 2016).

Apart from PfALBA1, cis-regulatory factors such as uORFs down-regulate the expression of dORFs (Kumar M, 2015). Frequent occurrence of uORFs in the transcripts of *P. falciparum* pose an additional layer of post-transcriptional gene regulation. One way to regulate the number of uORFs is by changing in the leader length of the transcripts which has been observed during different stages (Caro et al., 2014). For the remaining uORFs that still pose a hindrance to the ribosome, we propose that a mix of leaky scanning and reinitiation could regulate the expression of the dORF, not only during the IDC, but also during the numerous situations of stress that are faced by the parasite during multiplication and development in the human host.

Wide-spread use of reinitiation and leaky scanning has been observed during stress conditions in a multitude of eukaryotes (Andreev et al., 2015; von Arnim et al., 2014; Fijalkowska et al., 2017; Hinnebusch, 1993, 2005; Hinnebusch et al., 2016; Young et al., 2015; Zach et al., 2014). Seemingly, organisms facing stress undergo non-canonical translation to economise their energy and resource usage. Hence, investigating translation initiation factors during stress conditions in *P. falciparum* would provide insights into the role of multiple uORFs harboured by the parasite’s transcripts.

In addition to regulation of translation during stress, an assortment of non-canonical translation mechanisms including reinitiation, leaky scanning, internal ribosome entry, and ribosome shunting have been observed in viruses (de Breyne and Ohlmann, 2018; Ryabova et al., 2006). Given their small genome size, viruses have adopted multiple strategies to transcribe and translate their protein repertoire. In the case of *P. falciparum*, the constraint is not genome size, but the high AT content of the genome. Owing to this, mRNAs have numerous uORFs and uAUGs which the parasite’s translation machinery appears to handle by a widespread use of non-canonical translation mechanisms.

In conclusion, this is the first report that systematically delineates the features of uORFs that affect translation of the downstream gene in *P. falciparum*. Additionally, we show that a mix of reinitiation and leaky scanning mechanisms are employed in asexual stages of *P. falciparum* to translate the dORF in presence of upstream ORFs. Therefore, initiation factors such as PTEF, PfeIF1, PfeIF2 and PfeIF3 may be critically involved in translation and regulation of parasite proteins. This distinguishing feature of the *P. falciparum* cytoplasmic translation machinery has the potential to become a novel target for anti-malarial drugs.

## Supporting information

Supplementary datasheet 1

Supplementary datasheet 2

Supplementary datasheet 3

## Authors’ contributions

CK, MK, and SP conceived the study and designed the experimental set up. CK and MK carried out the experiments. CK and SP wrote the manuscript. All authors read and approved the final manuscript.

## Acknowledgements

This work was partially funded intramural funds from IIT Bombay. CK is supported by PhD Teaching-Assistant Fellowship from IIT Bombay. MK was supported by Research Fellowship from the University Grants Commission (UGC). We thank IIT Bombay hospital and staff for assisting in blood collection from the volunteers for parasite culturing.

